# Transcriptome-wide Identification of GlycoRNAs by Clier-seq Pipeline

**DOI:** 10.1101/2025.02.28.639800

**Authors:** Nannan Zhu, Yan-Lin Yang, Yuan-Tao Liu, Zheng-Zhou Lu, Yan Wang, Yi-Ling Luo, Ning Meng, Yan Yuan, Qian Zhong, Mu-Sheng Zeng

## Abstract

RNA molecules can undergo modification by N-glycans and be displayed on the cell surface. However, recent studies have focused primarily on N-glycan modifications on small RNAs less than 200 nt in length; the transcriptome-wide subtypes of glycosylated RNAs (glycoRNAs) remain poorly characterized. A validation system, which represents only a fraction of the total transcripts, has yet to be established to analyze glycoRNAs accurately. In this study, we aimed to comprehensively characterize transcriptome-wide global glycoRNAs and novel glycoRNA subtypes in both epithelial cells and B cells. Using metabolic labeling and density gradient centrifugation methods, we identified glycoRNAs predominantly below 2000 nt in both epithelial cells and B cells. We then developed the Clier-seq (Click chemistry-based enrichment of glycoRNAs sequencing) method to maximize the coverage of glycoRNAs (ranging from 50 to 2000 nt) and utilized the HISAT-StringTie-Ballgown pipeline to predict novel glycoRNA subtypes. We also established Clier-qPCR assays (click chemistry-based enrichment of glycoRNAs RT‒qPCR) to validate the specificity of the candidate glycoRNAs. We demonstrated that transfer RNAs (tRNAs), particularly tRNAs (Ser), tRNAs (Thr), tRNAs (Val), and tRNAs (lys), are the primary targets of glycosylation. Additionally, we found that vault RNAs (vtRNAs), specifically vtRNA2-1, are glycosylated. Furthermore, we discovered several novel glycosylated long noncoding RNAs ranging from 200–400 nt in length. Herein, we propose a standardized bioinformatics pipeline for glycoRNA research, enabling accurate and comprehensive identification of glycoRNAs throughout the transcriptome.

## INTRODUCTION

RNA plays a basic biological role in many cellular functions, including amino acid transport, protein translation, RNA transcription and modification (Dykstra et al., 2022). RNA undergoes various covalent modifications that alter its charge distribution, structure, catalytic activity, and interactions with proteins (Tyagi et al., 2023). For example, N6-methyladenosine (m(6)A), which is added by the METTL3/METTL14 methyltransferase complex, participates in RNA metabolism and regulates mRNA translation status and lifespan (He and He, 2021; Wang et al., 2014). tRNA molecules undergo multiple types of modifications, including methylations, monosaccharide modifications, and pseudouridine modifications, which regulate processes such as tRNA aminoacylation, ribosome recognition, and transport during protein synthesis (Suzuki, 2021). Recent studies have shown that RNA can also undergo N-glycosylation, promoting its localization on the cell membrane surface (Chai et al., 2023; Flynn et al., 2021; Ma et al., 2023; Nachtergaele and Krishnan, 2021; Perr et al., 2025). Many questions remain, such as which types of RNA molecules can be glycosylated and what specific functions glycoRNAs serve (Disney, 2021).

Recently, Flynn et al. established a metabolic labeling and click chemistry method to investigate glycoRNAs. The method involves the incorporation of an azide-modified sialic acid precursor, Ac4ManNAz, into glycoRNAs, followed by a click reaction with biotin probes. Using this technique, they demonstrated the presence of glycoRNAs in several cells and tissues and discovered that glycoRNAs were predominantly composed of small ncRNAs, specifically Y5 RNA (Flynn et al., 2021).

Their groundbreaking study revealed a novel RNA modification and significantly advanced our understanding of the role of RNA in extracellular biology. However, several limitations in glycoRNA sequencing and analysis need to be addressed. First, the glycosylation of small RNAs less than 200 nt in length was investigated, leaving the glycosylation of RNA above 200 nt unexplored. Second, it is unknown whether previously unannotated RNAs could undergo glycosylation. Additionally, the reads mapped to the reference genome accounted for only approximately 2% of the total, and the sequencing depth was biased toward Ac4ManNAz-enriched samples, potentially skewing the accurate identification of glycoRNAs (Flynn et al., 2021, Supplemental information). Finally, experimental controls are needed to validate candidate glycoRNAs and exclude nonspecific enrichment.

To address these issues, we made specific improvements in both the library preparation and analysis methods, leading to the development of the click chemistry-based enrichment of glyco<u>R</u>NAs (Clier-seq) pipeline for glycoRNA sequencing. The modifications include extending the RNA transcript length range from 0–200 nt to 0– 2000 nt via sucrose density gradient centrifugation. Additionally, we devised a novel glycoRNA library preparation method to increase the capture efficiency of long transcripts. For analysis, we employed the HISAT-StringTie-Ballgown pipeline to predict novel glycoRNAs. To validate the potential glycoRNAs, we established the Clier-qPCR (<u>cli</u>ck chemistry-based <u>e</u>nrichment of glycoRNAs <u>RT‒qPCR</u>) method, which incorporates experimental controls. Finally, we included a diverse range of cells and tissues for a comprehensive analysis of glycoRNAs.

By employing the Clier-seq pipeline, we achieved broad coverage and obtained a comprehensive overview of glycoRNAs at the whole-transcriptome level. We made several significant discoveries, including the confirmation of tRNA as a major subtype of glycoRNAs, the identification of glycosylated modified Vault RNA, and the discovery of previously unannotated long noncoding RNAs (lncRNAs) that undergo glycosylation.

## RESULTS

### 1. GlycoRNAs are present in epithelial cells and B cells

RNA molecules can be modified by N-glycans and presented on the cell surface, as well as in mouse liver and spleen tissues (Flynn et al., 2021). To detect glycoRNAs, we cultured the cells or tissues with Ac4ManNAz, allowing for the azide-modified sugars to be incorporated into the cellular polysaccharides. Following RNA extraction and purification, we performed a click chemistry reaction using biotin-conjugated probes to label glycoRNAs. The biotin-labeled glycoRNAs were subsequently detected through RNA blotting (Figure 1A).

**Figure 1.**
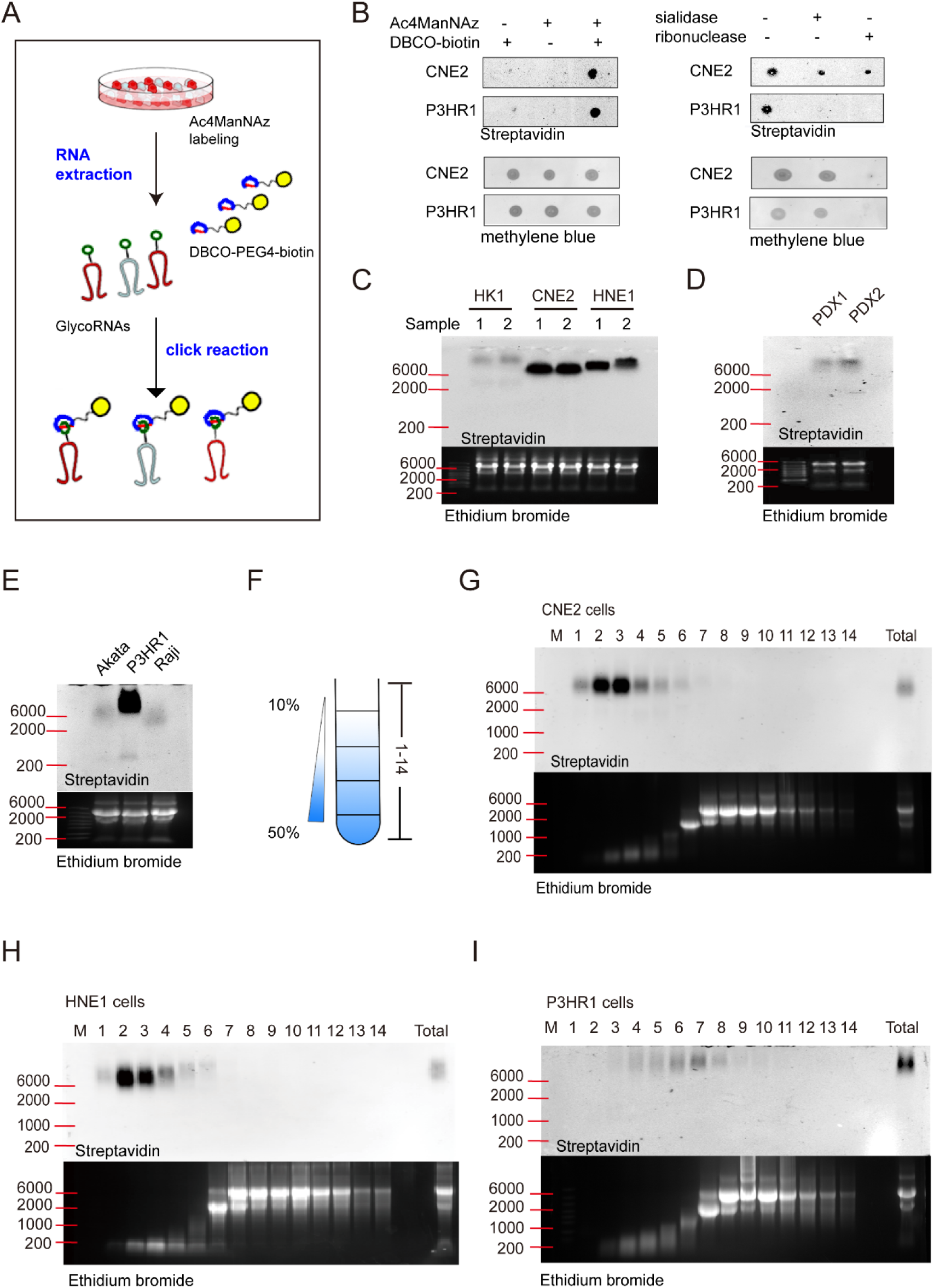
GlycoRNAs in epithelial and B cells. A. Schematic of glycoRNA metabolic labeling with Ac4ManNAz and the click reaction using DBCO-PEG4-biotin probes. (B) Dot blot of glycoRNAs in CNE2 and P3HR1 cells treated with Ac4ManNAz or DBCO-biotin and treated with sialidase or ribonuclease. C-E. GlycoRNA blotting of NPC cell lines (HK1, CNE2, and HNE1) (C), NPC-PDX tissues (D), and B lymphoma cell lines (P3HR1, Akata, and Raji) (E). F. Schematic of RNA sucrose density centrifugation, which involves fractionating RNA into 14 fractions through a 10%-50% sucrose gradient. G-I. GlycoRNA blotting of recovered RNA from distinct sucrose gradient fractions in CNE2 (G), HNE1 (H), and P3HR1 cells (I).

Figure 1 B shows that almost no spots were observed in controls lacking Ac4ManNAz metabolic labeling or when Ac4ManNAz labeling was performed without the subsequent click reaction. GlycoRNA spots were visible only when both labeling methods were combined, indicating the specificity of the labeling. In addition, treatment of the biotin-labeled RNA with sialidase or ribonuclease A significantly reduced the intensity of the glycoRNA spots, confirming that the RNA molecules were glycosylated (Figure 1B).

Using this metabolic labeling method, we detected glycoRNAs in various cell lines, including NPC cells (HK1, CNE2, and HNE1) (Huang et al., 1980; Kaitai et al., 1990; Zhu et al., 2022), NPC-PDX tissues, and B lymphoma cells (P3HR1, Akata, and Raji) (Kuhn-Hallek et al., 1995). We purified RNA from each cell type, labeled it with biotin-conjugated probes, and subjected it to formaldehyde gel electrophoresis. RNA blotting revealed the presence of glycoRNAs in both epithelial cells and B cells. The intensity and position of the glycoRNA bands varied among different cell types, indicating potential differences in the levels or states of RNA glycosylation (Figure 1C, D, E).

To investigate the size of glycoRNA transcripts, we purified total RNA and fractionated it through a sucrose density gradient ranging from 10% to 50%, resulting in the generation of 14 distinct fractions. Each fraction was subsequently subjected to RNA precipitation (Figure 1F). Following density gradient centrifugation, we performed RNA blotting to evaluate the levels of glycosylation in the recovered RNA transcripts (Figure 1G, H, I). GlycoRNA bands were observed in the 0-2000 nt range of RNA transcripts, which was consistent across the CNE2, P3HR1, and HNE1 cell lines.

Flynn’s supplementary data revealed that candidate glycoRNAs are primarily small ncRNAs in the range of 50-200 nt, such as snoRNAs, Y RNAs, and tRNAs (Flynn et al., 2021, Supplemental information). However, our sucrose density gradient centrifugation experiment revealed glycosylations on transcripts spanning from 0– 2000 nt. These findings suggest that glycoRNAs are not limited to small ncRNAs but may also exist on lncRNAs and mRNAs, indicating the potential existence of previously undiscovered glycoRNA subtypes.

These findings indicate a broader spectrum of glycoRNA types than previously known, prompting us to develop a comprehensive and accurate method for the identification of glycoRNAs within the 50–2000 nt range.

### 2. Clier-seq method for constructing glycoRNA sequencing libraries

To characterize transcriptome-wide global glycoRNAs and undiscovered subtypes of glycoRNAs in both epithelial and B cells, we established a Clier-seq (<u>Cli</u>ck chemistry-based enrichment of glyco<u>R</u>NAs <u>seq</u>uencing) analysis pipeline (Figure 2A, B, C). Clier-seq allows for the specific enrichment and sequencing of glycoRNAs, facilitating comprehensive analysis.

**Figure 2.**
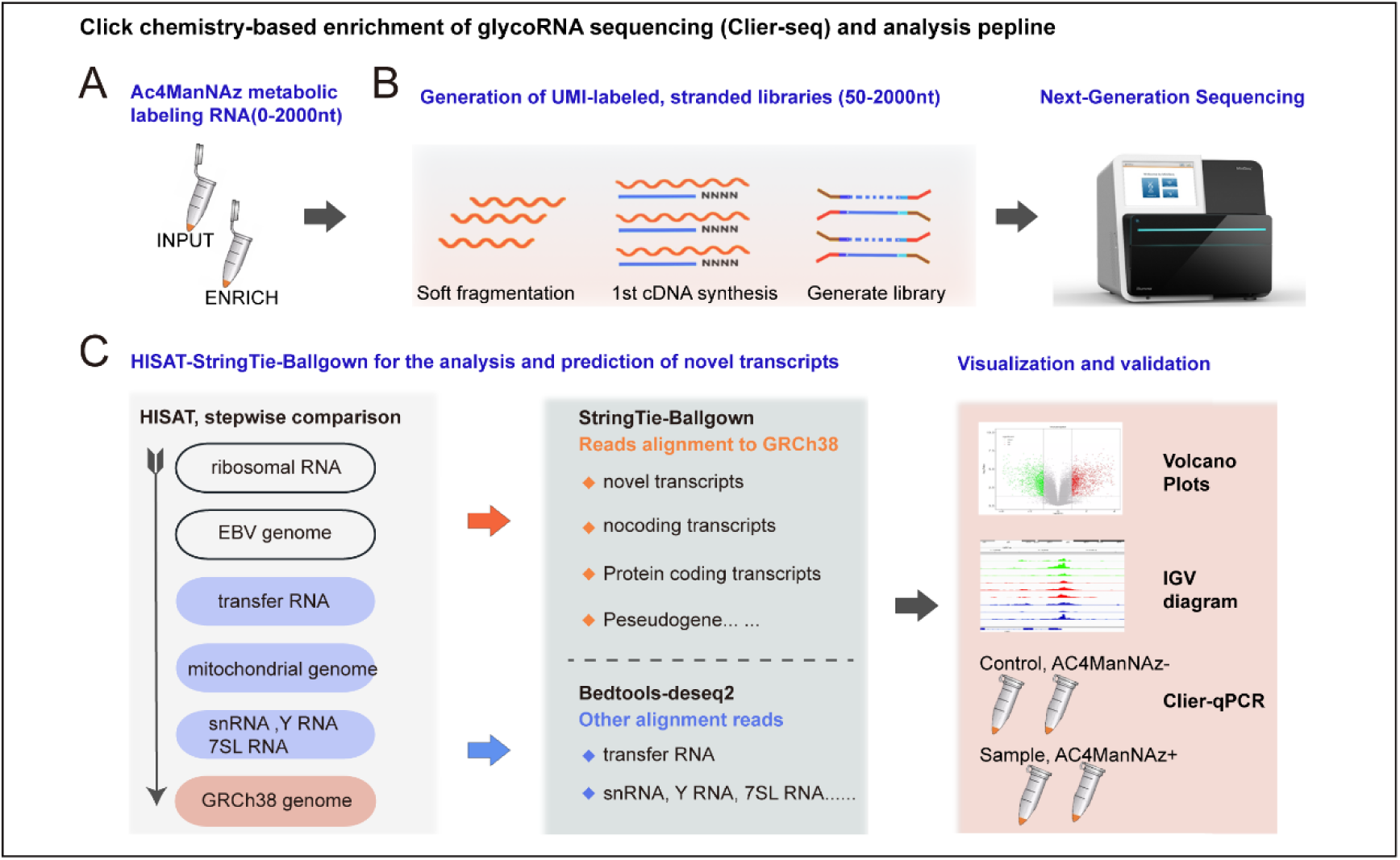
Schematic of the Clier-seq analysis pipeline. A. INPUT and ENRICH samples were prepared by purifying metabolically labeled RNA, treating it with DNase I and proteinase K, and collecting 0–2000 nt RNA transcripts. B. Strand-specific library preparation involved RNA fragmentation, cDNA synthesis, and ligation with UMI adaptors for sequencing. C. Sequence alignment via the HISAT-StringTie-Ballgown pipeline aligned processed reads to various genomes (EBV, mitochondrial, rRNA, tRNA, snRNA, Y RNA, and 7SL RNA) and the human genome reference GRCh38 to identify new glycoRNA transcripts.

We extracted and purified total RNA and then separated the RNA transcripts by size via sucrose density gradient centrifugation, focusing on those smaller than 2000 nt. The RNA was labeled with biotin via a click reaction. We retained an INPUT sample and enriched glycoRNAs via streptavidin magnetic beads, which we designated the ENRICH sample. Both samples were then used for library construction and analysis (Figure 2A and Figure S1A).

In Flynn’s study, the library construction protocol involved ligating an adaptor to the 3’ end of the RNA, synthesizing the first-strand cDNA, and then performing PCR to obtain the library sample. This method primarily detects small ncRNAs such as tRNA, snRNA, and Y RNA. The overall proportion of mapped reads was only approximately 20–30%, with human genomic reads accounting for approximately 2% of the total reads (Flynn et al., 2021, Supplemental information).

To address this issue, we developed a novel library construction method. In our protocol, we reverse transcribed first-strand cDNA via random primers and synthesized second-strand cDNA with dUTP for strand-specific library construction. We then ligated a UMI adaptor to the double-stranded cDNA and selected cDNA fragments by size via magnetic DNA beads before PCR amplification (Figure 2B and Figure S1B).

Sequencing was conducted on the Illumina NovaSeq PE150 platform, generating 6G of data for both the INPUT and ENRICH samples. Quality control analysis of the sequencing data revealed high quality, with Q20>95% and Q30>90%. The sequencing reads were then subjected to alignment against various references, including the EBV genome, rRNA, tRNA, mtRNAs, snRNA, Y RNA and 7SL RNA. Finally, the unaligned reads were aligned to the human genome reference GRCh38 (Figure 2C). The number of reads obtained was relatively consistent between the ENRICH and INPUT samples, with over 93% of the reads mapping to the target genome.

The RNA transcript lengths were mainly concentrated within the range of 50–2000 nt, covering key glycoRNA transcript lengths (Figure S1C). Our results revealed that the proportion of reads aligned to the human genome reference (GRCh38) increased after enrichment. For example, in the P3HR1 INPUT sample, 11.52% of the reads aligned to the GRCh38 genome reference, increasing to 16.90% after enrichment (Figure S1D).

This method effectively retained small RNA molecules, such as snRNAs, snoRNAs, and Y RNAs (Figure S1E, F), and increased the number of reads aligned to the human genome reference (GRCh38), outperforming Flynn’s protocol.

This standardized and efficient library construction process allows for rapid detection of glycoRNAs by completing the workflow from enrichment to library construction within a single day. We were able to retain the majority of previously identified candidate transcripts of glycoRNAs (small ncRNAs ranging in size from 50–200 nt). Moreover, we successfully captured a greater number of lncRNA and mRNA transcripts.

### 3. Identification of glycoRNAs via the HISAT-StringTie-Ballgown analysis pipeline

Our data analysis revealed that some enriched glycoRNA transcripts were in unannotated regions of the genome. Additionally, the original paper (Flynn et al., 2021) presented several issues: 1) a low alignment rate; 2) single-end alignment was incorrectly applied to paired-end data; and 3) differential analysis was conducted on raw read counts without normalization. These challenges prompted us to explore alternative methods for glycoRNA analysis.

To address these problems and comprehensively identify glycoRNAs, we employed HISAT2, StringTie, and Ballgown tools (Figure 2C), which enable genome alignment, transcript assembly, identification of novel splice variants, transcript abundance quantification, and differential expression analysis (Pertea et al., 2016). As part of this workflow, we systematically corrected the issues in the original paper, including adjusting the alignment of paired-end sequencing data, updating the human gene reference version, and using StringTie to normalize read counts (Figure S2A).

Our analysis focused on various epithelial and B-cell samples, including two NPC-PDX tissues, two nasopharyngeal epithelial cell lines (NP460 and NP69), three nasopharyngeal carcinoma cell lines (HK1, CNE2, and HNE1), and three B lymphoma cell lines (P3HR1, Akata, and Raji). We set INPUT or ENRICH as independent variables and used different cell samples and replicates as covariates, allowing for us to identify commonly enriched glycoRNAs across diverse cell types and replicates.

A volcano plot highlighted significantly enriched RNA transcripts in the ENRICH group, with the most enriched transcripts displayed in red on the positive fold-change side of the plot. Notably, many of these transcripts, such as MSTRG.7832, MSTRG.5930, and MSTRG.18836, were unannotated and predicted by StringTie, suggesting the presence of previously unidentified glycoRNAs (Figure 3A). The Integrative Genomics Viewer (IGV) plot revealed distinct expression peaks in different cell types while maintaining sequence consistency (Figure 3B). These findings underscore the importance of transcript assembly and prediction in glycoRNA identification across diverse samples.

**Figure 3.**
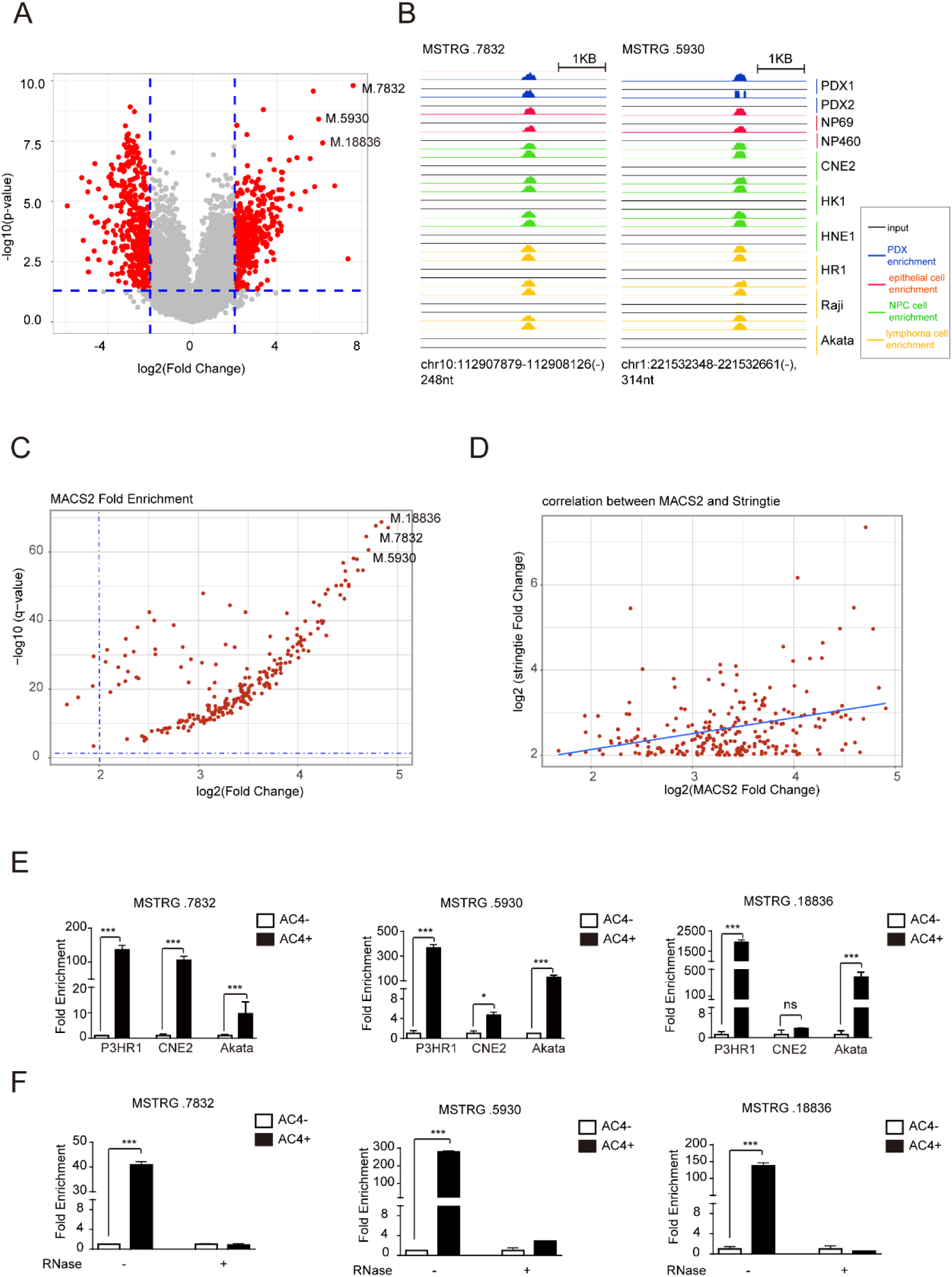
Discovery of glycoRNAs via the Clier-seq analysis pipeline. A. Volcano plot across all 16 cell lines and tissues showing Ac4ManNAz-enriched RNA transcripts, with highly enriched transcripts labeled. B. IGV diagram displaying representative highly enriched Ac4ManNAz RNA transcripts. C. GlycoRNA identification via the MACS tool. D. Correlation analysis between the MACS and Clier-seq results. E. Validation of enriched RNA transcripts in CNE2, Akata, and P3HR1 cells via Clier-qPCR. Biotin-labeled glycoRNAs were pulled down and analyzed by RT‒PCR. A control without Ac4ManNAz labeling (AC4-) was used as a reference, and Ac4ManNAz-labeled samples (AC4+) were compared (mean ± SD, n=3, p value by two-tailed unpaired t test). F. Cell surface RNAs from P3HR1 cells were either treated with RNase A or left untreated. The enriched RNA transcripts were validated via Clier-qPCR (means ± SDs, n=3; p values were determined via two-tailed unpaired t tests).

We also incorporated model-based analysis for ChIP-seq (MACS2), a tool traditionally designed for ChIP-seq, owing to its robust statistical model and peak-calling algorithm, which makes it applicable to RNA immunoprecipitation-based data. MACS2 enabled us to identify regions with enriched reads corresponding to glycosylation sites. The transcripts identified through MACS2 were consistent with those from the HISAT-StringTie-Ballgown pipeline, further validating our approach (Figure 3C, D).

### 4. Clier-qPCR method to validate the specificity of candidate glycoRNAs

One of the main experimental biases in glycoRNA metabolic labeling is the nonspecific binding of the biotin probe to certain RNAs. To address this, we conducted a Clier-qPCR (click chemistry-based enrichment of glycoRNAs <u>RT‒</u> <u>qPCR</u>) experiment.

We used an Ac4ManNAz-unlabeled sample as a negative control, which was processed identically to the experimental group but without Ac4ManNAz labeling. By calculating the enrichment-to-input ratio of the control sample, we determined the background enrichment level for a given transcript. The actual fold enrichment of the target transcript in the experimental group was then calculated by dividing the observed enrichment by the background enrichment ratio. This approach effectively excludes nonspecific signals generated during the biotin labeling and streptavidin bead enrichment process.

Using the Clier-qPCR method, we confirmed significant enrichment of MSTRG.7832, MSTRG.5930, and MSTRG.18836 in the P3HR1, Akata, and CNE2 cell lines, with particularly high fold enrichment values (Figure 3E). When P3HR1 cell surface RNA was treated with RNase A, the glycoRNA signal was completely abolished (Figure 3F). This experimental validation demonstrated the presence of glycosylations in the transcripts predicted by HISAT-StringTie.

Additionally, we examined other RNA transcripts that were highly enriched in the sequencing data, such as MSTRG.33163, MSTRG.35878 and MSTRG.39406. Consistent with the sequencing data, these transcripts also showed enrichment via Clier-qPCR, but their enrichment levels were not as prominent as those of MSTRG.7832 and MSTRG.5930. Notably, MSTRG.20675 was not significantly enriched according to Clier-qPCR (Figure S2B, C). The observed sequencing bias is likely due to the nonspecific binding of the biotin probe, highlighting the necessity of further screening and validation through Clier-qPCR.

In conclusion, our Clier-seq analysis pipeline integrates three modules to establish a standardized workflow for glycoRNA analysis. These three modules include 1) Clier-seq for library preparation and sequencing; 2) the HISAT-StringTie-Ballgown pipeline for bioinformatics analysis; and 3) Clier-qPCR for transcript detection and validation. This workflow enables comprehensive and accurate identification of glycoRNAs on a transcriptome-wide scale.

### 5. Transcriptome-wide analysis of RNA subtypes modified by glycosylation

We aimed to identify which types of RNA undergo glycosylation. To achieve this goal, we classified the RNA transcripts aligned to the genome reference (excluding tRNA, mtRNA, rRNA, snRNA, Y RNA, and 7SL RNA) into six categories on the basis of factors such as the NCBI gene type and status and GeneCards categories (Seal et al., 2020). The categories are as follows: 1) noncoding RNA (excluding snoRNA), 2) unannotated RNA transcripts predicted by StringTie, 3) pseudogene and other transcripts, 4) protein-coding RNA (excluding ribosomal protein mRNA), 5) ribosomal protein mRNA (RP mRNA), and 6) small nucleolar RNA (snoRNA).

Volcano plots from all 16 cell lines and tissues revealed that Ac4ManNAz-enriched transcripts predominantly belonged to the unannotated transcript category (Figure 4A), which is consistent with the results shown in Figure 3A. Additionally, several highly enriched transcripts in the ncRNA, pseudogene, and protein-coding categories were identified and validated through Clier-qPCR, with MSTRG.7832 serving as a positive control in the Clier-qPCR assay. Among these, vtRNA2-1 presented the highest enrichment ratio (Figure 4B).

**Figure 4.**
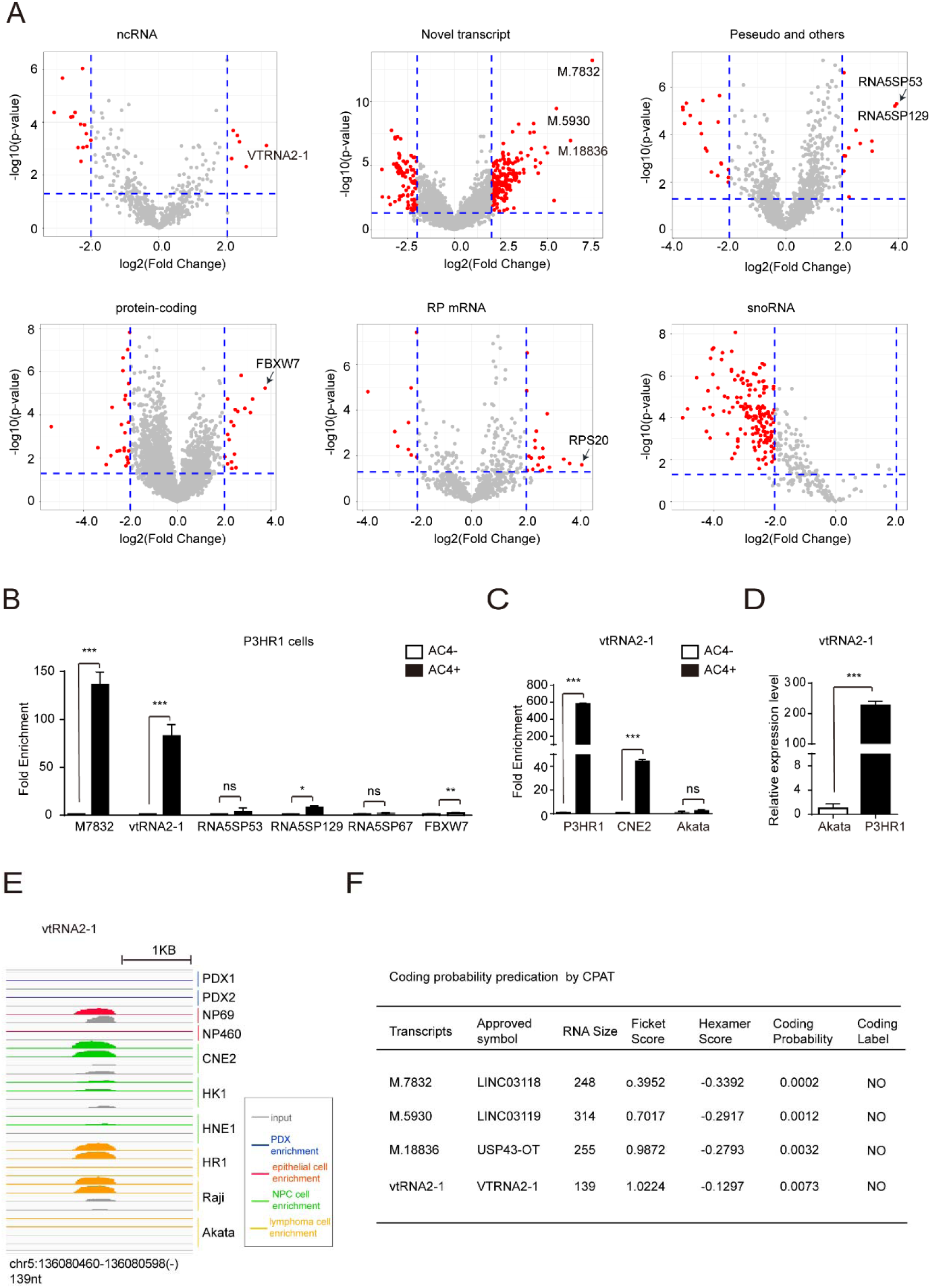
Identification of glycoRNAs in six RNA transcript categories. A. Volcano plots representing all 16 cell lines and tissues categorized into six RNA transcript types on the basis of NCBI Gene Classification and GeneCards categories. B. Validation of Ac4ManNAz-enriched RNA transcripts in P3HR1 cells via Clier-qPCR (means ± SDs, n=3; p values were determined via two-tailed unpaired t tests). C. Clier-qPCR analysis showing the fold enrichment of vtRNA2-1 in P3HR1, CNE2, and Akata cells (mean ± SD, n=3, p value from a two-tailed unpaired t test). D. RT‒ PCR analysis showing significant differences in the expression of vtRNA2-1 between Akata and P3HR1 cells (means ± SDs, n=3; p values were determined via two-tailed unpaired t tests). E. IGV diagram displaying vtRNA2-1 read alignment. F. Coding potential of glycoRNAs predicted via the RNA Coding Potential Assessment Tool (CPAT).

Clier-qPCR confirmed the high enrichment of vtRNA2-1 in P3HR1 and CNE2 cells but not in Akata cells (Figure 4B, C), likely due to its lower expression in Akata (1/230th of the level in P3HR1) (Figure 4D). Consistent with the Clier-qPCR results, the IGV diagram also revealed specific enrichment of vtRNA2-1 in P3HR1, Raji, and CNE2 cells (Figure 4E). vtRNA2-1, one of the four Vault RNAs (vtRNA1-1, vtRNA1-2, vtRNA1-3, vtRNA2-1), is a small ncRNA transcribed by polymerase III. These results suggest that vtRNAs could be candidates for glycosylated modification.

Three glycoRNA transcripts predicted from unannotated regions via StringTie (Figure 3A) were analyzed via the RNA Coding Potential Assessment Tool (CPAT). The analysis revealed that these transcripts had low coding potential and were longer than 200 nt (Figure 4F), classifying them as lncRNAs. We applied for gene symbols from the HUGO Gene Nomenclature Committee and named them linc03118, linc03119 and USP43-OT (Figure 4F). This discovery confirmed the presence of glycosylations in lncRNAs.

Overall, we found that partial transcripts of vtRNAs and lncRNAs exhibited glycosylation. There may be additional glycoRNA transcripts that were not identified in this study. Our results strongly suggest that glycoRNAs are specific to particular RNA types, primarily functioning as noncoding RNAs, with consistent transcript sequences across different cell types.

### 6. tRNA is a major subtype of glycosylated RNA

Through this study, we revealed that vtRNA and lncRNA can undergo glycosylations. Additionally, we performed enrichment analysis on RNA transcripts initially excluded from alignment, such as snRNA, Y RNA, 7SL RNA, tRNA, and mtRNA. Using ENRICH and INPUT as independent variables and considering all 16 cell lines and tissues as covariates, we screened the commonly expressed glycoRNAs via DESeq2 (Figure 2C).

Notably, we observed enrichment of tRNA transcripts, including tRNA-Ser-GCT-4-2, tRNA-Thr-AGT-1-1 and tRNA-Lys-TTT-7-1, as shown in volcano plots (Figure 5A). To eliminate biases from tRNA library preparation and sequencing, we validated the target transcripts via Clier-qPCR, confirming enrichment compared with the negative control sample (Figure 5B).

**Figure 5.**
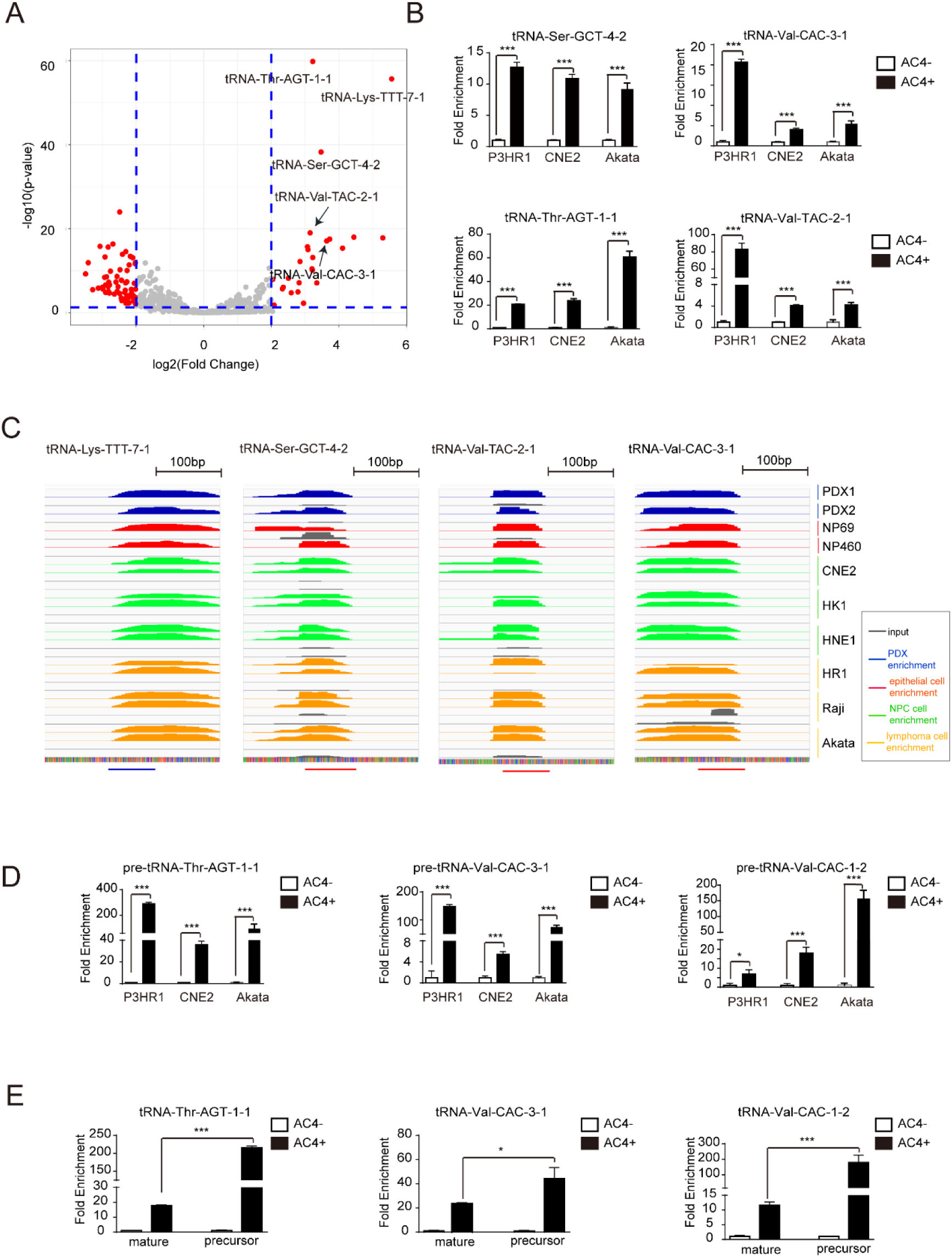
Analysis of tRNA glycosylation. A. Volcano plot of Ac4ManNAz-enriched tRNA transcripts. B. Clier-qPCR analysis showing the enrichment ratio of Ac4ManNAz-labeled tRNA in P3HR1, CNE2, and Akata cells (mean ± SD, n=3, p value by two-tailed unpaired t test). C. IGV diagram displaying read alignment of Ac4ManNAz-enriched tRNA transcripts. D. Validation of Ac4ManNAz-enriched pretRNA transcripts in P3HR1, CNE2, and Akata cells via Clier-qPCR (means ± SDs, n=3; p values were determined via two-tailed unpaired t tests). E. Comparison of the fold enrichment between tRNAs and pretRNAs via Clier-qPCR (means ± SDs, n=3; p values were determined via two-tailed unpaired t tests).

We further examined these enriched tRNA transcripts via the IGV, where homologous mature tRNAs are highlighted with blue (positive) or red (negative) lines. The analysis revealed that glycosylated tRNAs were enriched across all cell types. Some of these tRNAs presented longer 3’ ends than did mature tRNAs (Figure 5C). To investigate this further, we conducted Clier-qPCR assays on pretRNA. As shown in Figure 5D, transcripts such as pretRNA-Lys-TTT-7-1, pretRNA-Val-CAC-3-1, pretRNA-AGT-1-1 and pretRNA-Val-CAC-1-2 were significantly enriched in P3HR1, CNE2, and Akata cells. Additionally, pretRNAs presented significantly greater fold enrichment than mature tRNAs did (Figure 5E), suggesting that tRNA glycosylation may also occur in pretRNAs.

tRNAs are among the most abundantly modified RNA species. Recent studies have confirmed that glycosylation occurs at key tRNA modification sites. For example, Flynn identified the modified RNA base 3-(3-amino-3-carboxypropyl) uridine (acp3U) as a site of attachment of N-glycans in glycoRNAs (Xie et al., 2024). Moreover, vertebrate tRNAs for Tyr and Asp undergo queuosine (Q) modification, which is further glycosylated with galactose and mannose to generate galQ and manQ, respectively (Zhao et al., 2023). These glycosylation sites, such as acp3U and queuosine, are commonly found in tRNAs, highlighting their crucial role in glycosylations. Therefore, tRNAs represent an important class of glycosylated RNA.

## DISCUSSION

In summary, our research introduces a comprehensive method for analyzing glycoRNAs, addressing various limitations in earlier glycoRNA studies. By expanding transcript coverage to 0–2000 nt, we significantly enhanced the detection of long transcripts that earlier techniques, which were restricted to less than 200 nt, failed to capture. This extended coverage allowed for us to discover glycosylations in a wider variety of RNA types, including tRNAs, vtRNAs, and lncRNAs, and deepened our understanding of glycoRNA biology.

The HISAT-StringTie-Ballgown pipeline enabled us to investigate glycoRNA modifications in previously unannotated RNA transcripts. The application of Clier-qPCR, combined with an Ac4ManNAz-unlabeled control, helped eliminate nonspecific signals, increasing the accuracy of glycoRNA identification. Our methodology addresses issues associated with short transcript lengths, low alignment rates, and nonspecific biotin probe binding. This standardized pipeline is a valuable tool for a transcriptome-wide investigation of glycosylation.

In addition, our research offers a thorough analysis of glycosylation across different RNA types. This analysis has significant implications for future investigations into the localization and functions of glycoRNAs. Among the RNA types studied, tRNAs, vtRNAs, lncRNAs, and miRNAs (Figure S2D) exhibited significant glycosylation, whereas snoRNAs and mRNAs presented a lower probability of glycosylation.

For example, we confirmed that tRNAs constitute a major class of glycoRNAs, identifying glycosylation-modified tRNA subtypes such as tRNAs (Ser), tRNAs (Thr), tRNAs (Lys), and tRNAs (Val). Our findings imply that glycosylation may take place during tRNA maturation, similar to other modifications, such as m1A58 (Li et al., 2022) and inosine (Torres et al., 2015). Abnormal tRNA modifications are strongly associated with diseases such as cancer and neurological disorders (Suzuki et al., 2011), making the investigation of tRNA glycosylation essential for a deeper understanding of disease mechanisms.

Moreover, we identified glycosylated vault RNAs, particularly vtRNA2-1, which play roles in cellular functions such as drug resistance, apoptosis, and viral replication (Amort et al., 2015; Büscher et al., 2020). Gaining insight into vault RNA glycosylation is essential for clarifying its functional roles and connection to diseases such as cancer (Treppendahl et al., 2012; Yu et al., 2020). We also identified three glycosylation-modified lncRNAs, namely, LINC03118, LINC03119, and USP43-OT. LINC03118 and LINC03119 are located between genes, whereas USP43-OT partially overlaps with the exons and introns of USP43, a histone deubiquitinase linked to tumor progression (He et al., 2018; Xue et al., 2022; Ye et al., 2021), suggesting that glycosylation might influence the function of these lncRNAs and affect disease progression.

Additionally, our research examined glycosylation in other RNA types, such as snoRNAs; however, low enrichment levels, as indicated by the volcano plot (Figure 4A), IGV diagram (Figure S3A), and Clier-qPCR results (Figure S3B), imply that snoRNAs such as SNORD10 and SNORA7B are unlikely to undergo glycosylation or that such modifications may be specific to certain cell types. Similarly, the analysis of mRNA glycosylation revealed that ribosomal protein (RP) mRNAs, such as RPS20, RPL21, and RPS24, exhibited low enrichment (Figure S3A, C), suggesting that glycosylation of RP mRNAs may be rare or restricted to particular cellular contexts.

Despite the advancements made, our study has several limitations. It focuses on larger glycoRNAs (50–2000 nt), resulting in lower detection efficiency for transcripts smaller than 50 nt, such as miRNAs and tiRNAs. In addition, our techniques do not fully account for the potential effects of glycosylation on library preparation and sequencing. Emerging methods such as modification-induced misincorporation tRNA sequencing (mim-tRNAseq) (Behrens et al., 2021) or nucleoside analysis through liquid chromatography‒mass spectrometry (Suzuki, 2021) may address these challenges and increase the accuracy of glycoRNA detection in future research.

Furthermore, our study focused on commonly occurring glycoRNAs within a limited array of cell lines and patient-derived xenografts (PDXs). We did not investigate cell- or disease-specific glycoRNAs, although our data indicated discrepancies in glycosylation among different cell types. Certain glycoRNAs may be restricted to specific cell types rather than being universally present. A deeper understanding of the expression and modification of glycoRNAs in various pathological contexts, along with the identification of cell- or disease-specific glycoRNAs, could greatly advance their use as biomarkers for disease diagnosis, prognosis, and treatment.

## MATERIALS AND METHODS

### Cell culture

The human normal nasopharyngeal cell lines NP69 and NP460 were generated by SW Tsao’s group at the University of Hong Kong. NP69 cells were maintained in keratinocyte-free media supplemented with EGF and pituitary serum (Invitrogen, Carlsbad, CA, USA), while NP460 cells were cultured in a medium comprising a 1:1 mixture of EpiLifeTM medium supplemented with 1% EpiLife®-defined growth supplement (Life Technologies, Carlsbad, CA, USA) and keratinocyte serum-free medium enriched with 0.2% defined keratinocyte growth supplement (Life Technologies, Carlsbad, CA, USA). Nasopharyngeal carcinoma cells (HK1, CNE2, HNE1) and B lymphoma cells (P3HR1, Akata, Raji) were cultured in RPMI-1640 media supplemented with glutamine, 10% fetal bovine serum (FBS) and 1% penicillin/streptomycin (P/S). The cells were grown in a humidified 5% CO2 incubator at 37 °C.

### Metabolic chemical reporters

N-azidoacetylmannosamine-tetraacylated (Ac4ManNAz, Click Chemistry Tools) was prepared at a concentration of 100 mM in sterile dimethyl sulfoxide (DMSO) and stored at −80 °C. For the metabolic labeling experiments, the cells were exposed to Ac4ManNAz at a final concentration of 50 µM for 48–72 hours, depending on the cell or tissue passage/growth rate, meaning that the labeling time corresponded to one cell generation.

### RNA extraction, DNase I treatment and proteinase K treatment

We employed the standard TRIzol reagent (Thermo Fisher Scientific) extraction method to isolate RNA. The aqueous phase was precipitated with 2x volumes of 100% ethanol (EtOH) and further purified via Zymo RNA Clean & Concentrator columns, which strictly adhered to the manufacturer’s instructions. Typically, RNA samples were eluted in 90 µl of RNase-free water, and their concentrations were determined via a Nanodrop (Thermo Fisher Scientific). For additional RNA purification, to digest the DNA, we added 2 µl of recombinant DNase I (Takara), 10 µl of 10X DNase I Buffer, and 1 µl of recombinant RNase Inhibitor (Takara) to 40-100 µg RNA and incubated it at 37 °C for 20 min. Protein digestion was performed by adding Proteinase K (PK, Thermo Fisher Scientific) to the RNA sample at a final concentration of 200 µg/ml, followed by incubation at 50 °C for 15 minutes. After PK digestion, the RNA was further purified via Zymo RNA Clean & Concentrator columns.

### Copper-free click conjugation to RNA

The RNA samples were mixed with an equal volume of dye-free gel loading buffer II (df-GLBII), containing 95% formamide, 18 mM EDTA, and 0.025% SDS, and incubated at 70 °C until denaturation. A 10-mM stock solution of dibenzocyclooctyne-PEG4-biotin (DBCO-PEG4-biotin, Sigma) was added to the denatured RNA, yielding a final concentration of 500 µM. The conjugation reaction was conducted at 55 °C for 10 minutes.

### Enzymatic treatment of RNA samples

For RNA digestion, we added 1 μl of ribonuclease A (Takara) to 10 μl of the RNA sample and incubated it at 37 °C for 1 h. To digest the glycans, we added 2 μl of 10X GlycoBuffer and 1 μl of α2-3,6,8,9-neuraminidase A (NEB) and incubated the mixture at 37 °C for 1 hour.

### Gel electrophoresis, blotting, and imaging

The RNA, which was purified and conjugated with DBCO-PEG4-biotin reagent as described earlier, was resuspended in 15 µl of nuclease-free water. In our setup, 25 µg of RNA from cell samples was loaded onto a gel. The RNA was mixed with an equal volume of 2x RNA gel loading dye (Thermo Fisher Scientific), denatured at 55 °C for 10 minutes, and then cooled on ice for 3 minutes. The samples were loaded onto a 1% agarose‒formaldehyde denaturing gel (1% agarose, 1× MOPS, and 6% formaldehyde) and subjected to electrophoresis at 60 V for 60 minutes. The RNA bands were visualized via a UV gel imager. For RNA transfer, a capillary transfer protocol was employed with a 0.45-µm nitrocellulose membrane (NC, Millipore). The RNA was cross-linked to the NC membrane via UV-C light (0.125 J/cm2), and the membrane was blocked with 5% BSA for 45 minutes at room temperature. The blocking buffer was prepared using PBST (1X PBS, 0.1% Tween-20). Following blocking, the membrane was stained with streptavidin-IR800 (LI-COR Biosciences) diluted to 1:2000 in blocking buffer for 30 minutes. The membranes were washed with PBST and visualized via LI-COR software.

### Dot blotting

The RNA, previously conjugated with DBCO-PEG4-biotin, was diluted to a final concentration of 500 ng/μl. After the RNA was incubated at 95 °C for 3 minutes to disrupt secondary structures, the samples were cooled on ice. One microliter of RNA (500 ng) was carefully spotted onto nitrocellulose, which was prepared and cross-linked as described earlier. After cross-linking, the membrane was immersed in a 0.2% methylene blue solution for 10 minutes and then washed with distilled water. A photo was recorded under white light to document the amount of loaded RNA as a control. Subsequently, further blocking steps were performed. Imaging was conducted via LI-COR software.

### Sucrose gradient fractionation of RNA

RNA treated with proteinase K (PK) was subjected to sedimentation through a discontinuous sucrose gradient on the basis of McConkey’s method (McConkey, 1967). We lyophilized 300 μg of total RNA and reconstituted it in 300 µl of buffer containing 50 mM NaCl and 100 mM sodium acetate (pH 5.5). To prepare the sucrose gradient, 1×5.5 ml polypropylene tubes (Beckman) were used. Gradients were formed in 5.5-ml tubes with 50%, 40%, 30%, 20%, and 10% sucrose solutions. This sequential procedure ensured the gradual formation of the gradient, with careful attention to prevent intermixing of solutions during the process. The reconstituted RNA was layered on top and centrifuged using an SW55 Ti rotor at 80000x g for 18 h in a Beckman Coulter L-100XP Ultracentrifuge, all of which were maintained at 4 °C. We sequentially transferred 360 µl of the sample from top to bottom into the collection tube. To assess the RNA distribution, 10 µl aliquots from each tube were loaded onto a 1% agarose gel and subjected to electrophoresis at 120 V for 20 minutes. After this, the remaining 350 µl of each sample was supplemented with a 1/10 volume of 3 M sodium acetate (pH=5.2) and 2.5 volumes of 100% EtOH. The mixture was precipitated at −80 °C for a minimum of 2 hours in preparation for subsequent sequencing.

### Enrichment of Ac4ManNAz-labeled RNA and sequencing

We extracted total RNA from NP69, NP460, HK1, CNE2, HNE1, Raji, Akata, and P3HR1 cells that had been previously labeled with Ac4ManNAz following the purification process described earlier. For increased reliability, biological replicates were generated by utilizing cells from distinct passages in the sequencing experiments.

For initial enrichment, sucrose gradient fractionation was performed, and RNA fractions less than 2000 nucleotides (nt) were collected after centrifugation and precipitated with 100% EtOH. In the second round of enrichment, a selective affinity approach using streptavidin beads was employed. We reserved 150 ng of the sorted RNA as the INPUT sample, whereas 25 µg of the sorted RNA was used as the ENRICH sample for enrichment via streptavidin magnetic beads.

To construct sequencing libraries, we prepared chain-specific libraries with the Universal V6 RNA-seq Library Prep Kit (VAHTS), and then starting cDNA synthesis with the Universal V6 RNA-seq Library Prep Kit (VAHTS) via Dual UMI Adapters from Dual UMI UDI Adapters Set 1 for Illumina (VAHTS) for ligation. Ligated RNA was purified with VAHTS DNA Clean Beads.

The small RNA-seq libraries were generated via the VAHTS Small RNA Library Prep Kit (Vazyme Biotech, NR801) according to the manufacturer’s protocol. Libraries were sequenced on the Illumina NovaSeq 3000/4000 platform to produce paired-end 150-bp reads.

### cDNA synthesis for RT‒qPCR analysis

Total RNA from Ac4ManNAz-labeled CNE2 and P3HR1 cells was extracted, purified, and clicked to DBCO-PEG4-biotin. Thirty micrograms of biotinylated total RNA serving as the ENRICH sample was diluted in biotin wash buffer for enrichment, and 300 ng of each sample before enrichment was saved as the INPUT sample.

cDNA synthesis was initiated following the instructions of the RevertAid First Strand cDNA Synthesis Kit (Thermo Fisher Scientific) in a 20 µl reaction mixture

### SYBR Green RT‒qPCR

Each PCR was performed in a 10 μl volume of 5 μl of 2×RealStar Green Fast Mixture (GenStar), 1 μl of primer mixture at 0.5 μM (each), 2 μl of cDNA template and 2 μl of nuclease-free water. PCR was carried out via a standard protocol.

### Calculation of fold enrichment

The INPUT samples labeled with Ac4ManNAz were denoted as INPUT_AC4+_, and the corresponding ENRICH samples were designated as ENRICH_AC4+_. Conversely, the Ac4ManNAz unlabeled control samples are referred to as INPUT_AC4−_ and ENRICH_AC4−_. Following the CHIP‒qPCR analysis methodology (Solomon et al., 2021), some modifications were made:

ΔCt [normalized ENRICH] = (Ct [ENRICH] - (Ct [INPUT] -Log2 (Input Dilution Factor))

ΔΔCt [Clier] = ΔCt [normalized ENRICH_AC4+_] - ΔCt [ normalized ENRICH_AC4−_] Fold Enrichment = 2^ (-ΔΔCt [Clier])

### Data analysis

The raw sequencing data were processed with fastp (Chen et al., 2018) to remove adapters. Next, HISAT2 was used to map the reads to a reference. In detail, the reads were aligned to the Epstein–Barr virus (EBV) reference genome, repetitive sequences, ribosomal RNA sequences, tRNA sequences, the mitochondrial genome, and the reference genome (GRCh38) in a stepwise manner. Picard was used for deduplication, and a gencore (Chen et al., 2019) was used if it contained UMI. Like the approach used by Flynn et al. (Flynn et al., 2021), we used bedtools to count reads of manually selected sequences and performed differential expression analysis via deseq2 (Love et al., 2014; Quinlan and Hall, 2010). For the reads that aligned to the genome, we used stringtie (Pertea et al., 2015) to assemble new transcriptions and quantify them. For these transcripts, we performed differential analysis via Ballgown (Frazee et al., 2015). Specifically, for a single cell line, we used two biological replicates and set the sample number as a covariate to identify enriched transcripts that are common to all cell lines. To increase the reliability of our results, we used MACS2 (Zhang et al., 2008) for peak-based enrichment analysis and combined the results from StringTie and MACS2 via bedtools. Statistical analysis was performed via R, and plots were generated via ggplot2. Source code for bioinformatics analysis: https://github.com/bigtaotao/Clier-seq.git

## Author contributions

Experimental design: N.Z. Experimental performance: N.Z., Y-L.Y., Z-Z.L., M.N. Data analysis: Y-T.L. Provision of materials (NPC-PDX tissue samples): Y-L.L. Project guidance: M-S.Z., Q.Z., Y.W., Y.Y. Manuscript writing and revision: N.Z., Y-L.Y., M-S.Z.

## Compliance and ethics

The authors declare that they have no competing interests. The research involving animals was reviewed and approved by the Institutional Animal Care and Use Committee (IACUC) at Sun Yat-Sen University, approval no. SYSU-IACUC-2018-000193. The experiments were conducted in accordance with the institutional guidelines for the care of laboratory animals, as published by the Ministry of Science and Technology of the People’s Republic of China.

## Acknowledgments

This study was supported by the National Key Research and Development Program of China (2022YFC3400900), the National Natural Science Foundation of China (81830090 and 82321003, 82202510, U22A20322 and 82030046), the Program for Guangdong Introducing Innovative and Entrepreneurial Teams (2019BT02Y198), the Guangdong Science and Technology Department (2020B1212030004), and the Sun Yat-sen University Talent Program (22yklj08).

We would like to express our gratitude to ChatGPT for its assistance in correcting and improving our written English. We would like to express our gratitude to Hu Ti-Rong for setting up the bioinformatics analysis system and providing guidance on bioinformatics analysis methods.

---

**Figure S1.**
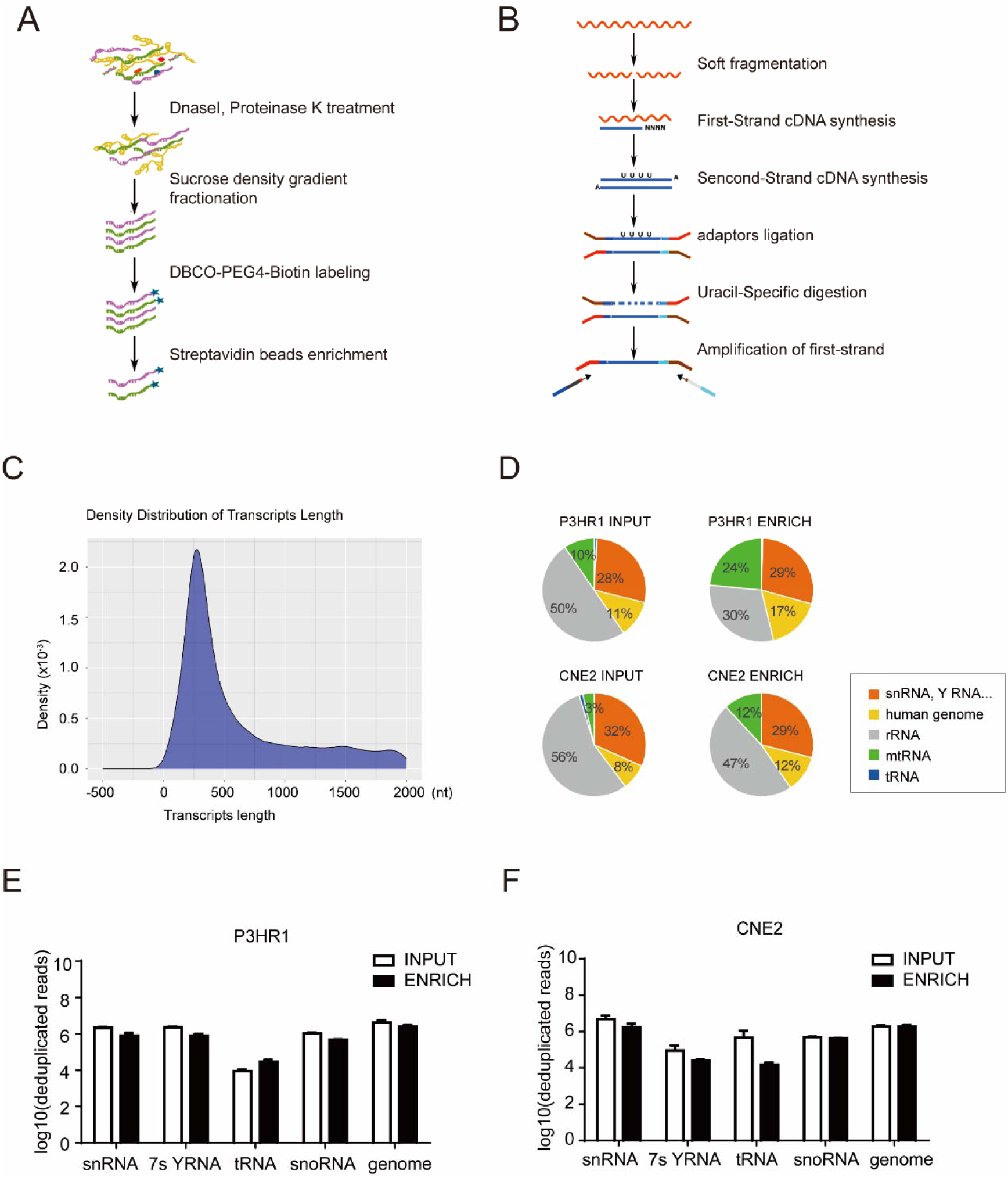
Establishment of the Clier-seq method for GlycoRNAs. A. Purification and enrichment of 0–2000 nt Ac4ManNAz-labeled RNA transcripts. B. Strand-specific library preparation. C. Histogram of RNA transcript lengths, ranging from 50–2000 nt. D. Pie chart of RNA types aligned to the GRCh38 genome reference. E, F. Deduplicated read counts of various RNA types from the INPUT and ENRICH libraries in P3HR1 and CNE2 cells.

**Figure S2.**
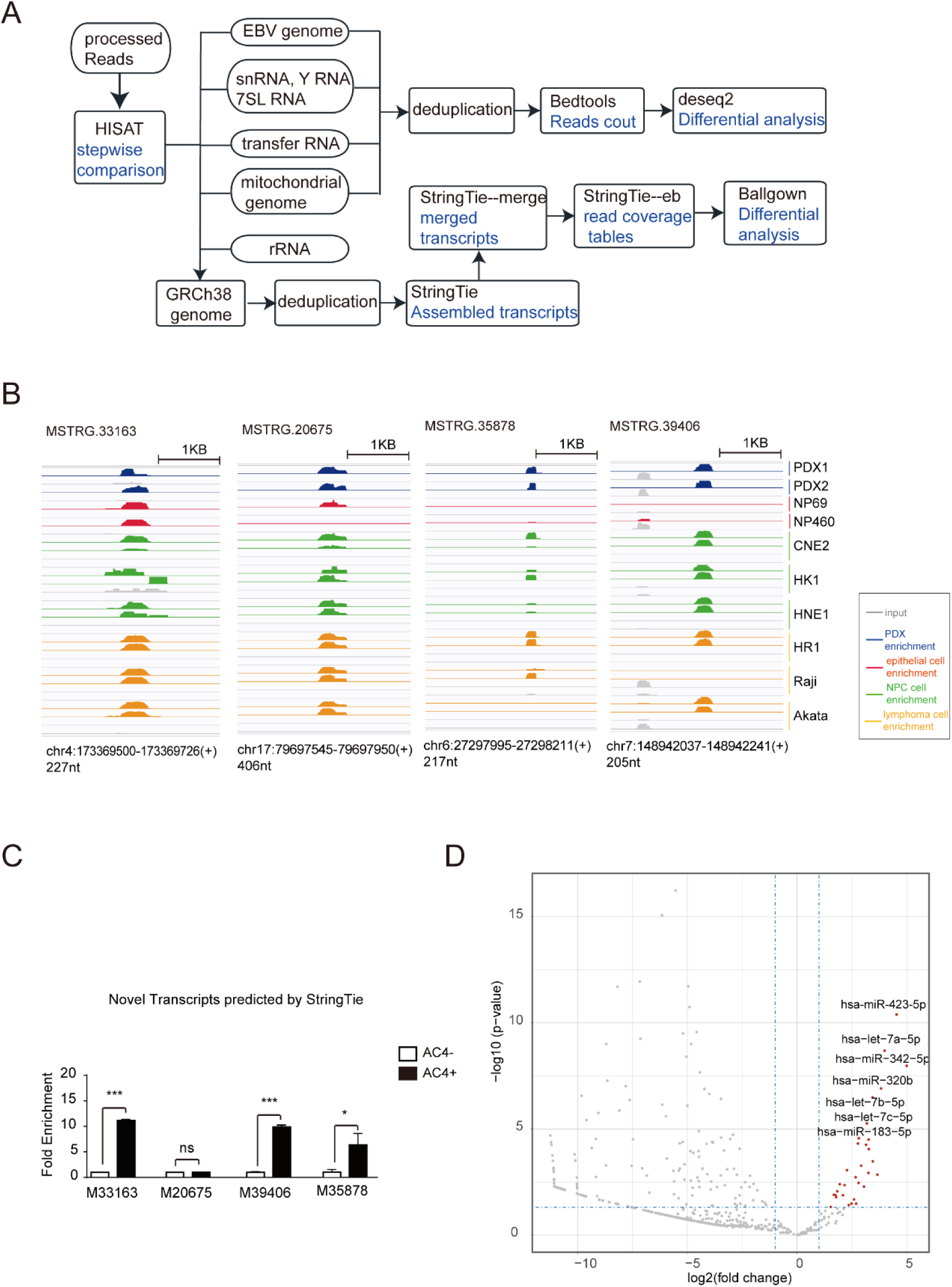
Analyzing glycosylation via the HISAT-StringTie-Ballgown pipeline. A. Schematic of the HISAT-StringTie-Ballgown analysis pipeline. B. IGV diagrams showing Ac4ManNAz-enriched RNA transcripts. C. Validation of Ac4ManNAz-enriched RNA transcripts in P3HR1 cells via Clier-qPCR (means ± SDs, n=3; p values were determined via two-tailed unpaired t tests). D. Volcano plot of Ac4ManNAz-enriched miRNA transcripts.

**Figure S3.**
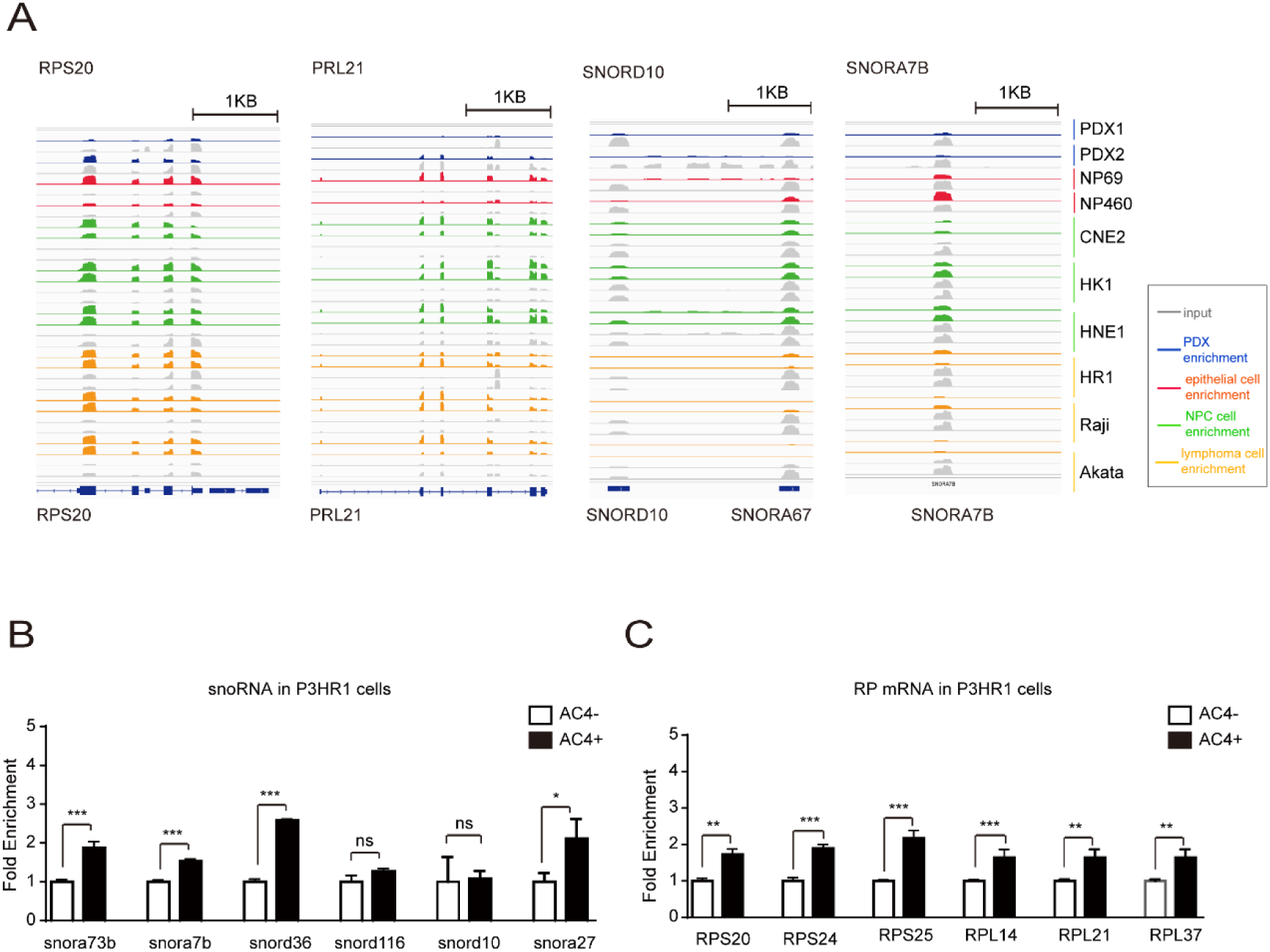
Glycosylation modifications in different RNA categories. A. IGV diagram showing read alignment of RPS20, RPL21, SNORD10, and SNORA7B. B, C. Validation of Ac4ManNAz-enriched snoRNAs (B) and RP mRNAs (C) in P3HR1 cells via Clier-qPCR (means ± SDs, n=3; p values were determined via two-tailed unpaired t tests).

## Notes

### Competing Interest Statement

The authors have declared no competing interest.

